# Fuzzifier*: Robust and Sensitive Multi-omics Data Analysis

**DOI:** 10.64898/2026.02.06.701074

**Authors:** Felix Offensperger, Chuqiao Pan, Evi Sinn, Ralf Zimmer

**Affiliations:** Institute of Bioinformatics, Department of Informatics, LMU Munich, Germany; Leibniz Institute for Food Systems Biology, Freising, Germany

**Keywords:** Uncertain Data Modeling, Categorization, Fuzzification, Hypothesis testing, Data Analysis, Transcriptomics, Cancer miRNAs

## Abstract

**Motivation:** Categorization is an important means for interpreting data and drawing conclusions. Often, the derived categories provide evidence for diagnostic or even therapeutic approaches. The standard pipelines for differential analysis of multi-omic high-throughput, and in particular single-cell data, yield (ranked) lists of possibly differential features after applying appropriate effect sizes or significance thresholds of computed *p*-value and/or foldchange.

**Results:** We propose the *Fuzzifier^*^* pipeline for the differential analysis of any type of high-throughput data, either raw input data or fold-change data of groups of a (small or large) number of replicates. In *Fuzzifier^*^*, categorization can be applied to any step of the analysis pipeline according to custom-designed fuzzy concepts (*Fuzzifier*). Thus, any (fuzzified) analysis option corresponds to a path in a ‘commutative diagram’ specifying the *Fuzzifier^*^* pipeline. *Fuzzifier^*^* computes a user-defined set of paths and presents an overview of the results, thereby identifying both highly reliable (consensus) and sensitive (path-specific) features. *Fuzzifier^*^* is a method that can be applied to any analysis pipeline to obtain different views on the data and yield more reliable results. This is demonstrated by the identification of context-specific miRNAs for individual cancer types from TCGA data. *Fuzzifier^*^* could both validate known cancer-specific miRNAs and identify novel candidates. In comparison to statistical tests, *Fuzzifier^*^* focuses on value distributions of tumor and normal samples as well as paired foldchange distributions and, thus, identifies condition-specific features from a relatively small number of replicates.

**Availability and Implementation:** https://github.com/zimmerlab/fuzzifier

## 1 Introduction

Among the tools we use to impose order on a complicated (but by no means unstructured) world, classification—that is, the division of things into categories based on perceived similarities—must be considered the most general and comprehensive of all.

Stephen Jay Gould, Questioning the Millennium, 1999.

Decades of high-throughput sequencing and other omics technologies have transformed biological research, providing unprecedented insights into complex biological systems. However, the interpretation of such data remains challenging due to inherent uncertainty, technical bias, and measurement imprecision. These issues of deriving robust, reliable and trustworthy conclusions via statistical (or bioinformatical analysis) have intensively described and discussed in the context of the so called ‘reproducibility crisis’ of biomedical sciences [8], and also in other sciences such as psychology [20], and, recently in ecological sciences [5]. Conventional analytical pipelines often rely on rigid mathematical formulations or arbitrarily chosen thresholds, such as fixed foldchange or significance cutoffs. While these approaches are simple to apply, they frequently fail to capture the gradual and context-dependent behavior of biological data.

In many omics analyses, a large number of quantitative values must be aggregated, summarized, or discretized. These operations often collapse continuous variation into binary categories, leading to decisions based on a single oversimplified distribution or statistic. As a result, subtle but biologically meaningful changes may be ignored, and extremes driven by preprocessing, normalization, and modeling choices are overemphasized. This issue becomes increasingly severe as experimental designs grow more complex and datasets become higher-dimensional, where traditional formulations struggle to produce interpretable and robust results.

Fuzzy logic offers a principled alternative to address these challenges. Unlike classical binary logic, in which propositions are either true or false, fuzzy logic allows variables to assume degrees of belief on a continuous scale between 0 and 1 [27, 17]. This formalism enables uncertainty and imprecision to be explicitly represented rather than ignored. Biological measurements such as gene expression levels, protein abundances, or metabolite concentrations are inherently continuous and noisy, which makes fuzzy logic particularly suitable for their interpretation [9, 25, 16, 18]. Although, there are appropriate methods for necessary data normalization [7, 28], the specific normalization can heavily influence the computation of effect sizes, associated significances and, thus, the conclusions. Fuzzification can help to avoid speculative results and add additional confidence in the conclusions.

In omics data analysis, key decisions, for example, whether a gene is expressed, differentially expressed, or mutated, are often treated as binary outcomes derived from these paradigms. Small changes in thresholds or modeling assumptions can produce substantially different feature sets, complicating biological interpretation and reproducibility. Fuzzy sets provide a unified framework for expressing such decisions, replacing hard assignments with degrees of membership that reflect both effect size and confidence, which together form a ‘degree of belief’.

Probably the most commonly used categorization during the analysis of high-throughput data expression data is the assignment of genes to meaningful linguistic categories such as ‘*upregulated*’, ‘*down-regulated*’, or ‘*unchanged*’. Differential expression analysis has become the de facto framework for addressing this task and is predominantly implemented through hypothesis-testing–based methods, including widely used tools such as DESeq2 [10], edgeR [19], and single-cell pipelines in Scanpy [26] and Seurat [21]. These approaches formulate gene-wise null hypotheses and rely on *p*-values, combined with multiple-testing correction, to derive categorical decisions.

However, the reliance on null hypothesis significance testing in this context has been increasingly criticized. Both practical and conceptual limitations have been highlighted, including the sensitivity of *p*-values to sample size, the frequent violation of model assumptions, and the conflation of statistical significance with biological relevance [24, 12]. As a consequence, statistically significant results may correspond to negligible effect sizes, while biologically relevant changes may be dismissed due to insufficient power.

An alternative perspective focuses directly on effect size, rather than hypothesis rejection [6]. When gene-wise foldchange distributions are available, the magnitude and direction of expression changes can be quantified and interpreted without invoking a null hypothesis. In this view, differential expression becomes a problem of estimating and contextualizing effect sizes under uncertainty, rather than testing for the existence of an effect. This paradigm has been advocated in multiple settings as a more interpretable and biologically aligned approach to expression analysis [14].

A complementary abstraction arises from considering expression levels themselves. Fold-changes, by definition, describe transitions between expression states rather than absolute quantities. Defining linguistic expression levels (e.g., ‘*low*’, ‘*medium*’, ‘*high*’) and interpreting foldchanges as transitions between such levels provides a natural and intuitive representation of differential expression. This perspective aligns closely with human reasoning about biological systems and lends itself to rule-based and fuzzy-logic formulations.

Despite its conceptual suitability, fuzzy logic has seen limited adoption in mainstream omics analysis. Existing applications have largely focused on clustering, classification, or rule-based modeling, while core analytical steps such as normalization and differential feature detection remain dominated by crisp methods. A general framework for integrating fuzzy logic across omics analysis workflows is still lacking.

In this work, we present a systematic approach for incorporating fuzzy sets and fuzzification into omics data analysis. While we use high-throughput sequencing of miRNA data for illustration, the framework is applicable to any omics pipeline. We demonstrate that the interpretation of data can be applied to multiple steps of the data analysis, leading to a more robust analytical framework, where the design decision for the analysis is questioned. By preserving uncertainty in the categorization throughout the analysis pipeline, this approach aims to improve interpretability, robustness, and biological relevance.

## 2 Methods

To integrate fuzzy logic into omics data analysis, we introduce a general fuzzification framework (*Fuzzifier* ) designed to categorize the data together with modeling uncertainty in this decision, reduce reliance on arbitrary thresholds, and support downstream analyses such as normalization and differential feature detection. The framework transforms quantitative measurements into fuzzy representations while preserving interpretability and mathematical consistency.

### 2.1 Fuzzification

Fuzzification is the process of mapping numerical measurements to degrees of membership in linguistically interpretable categories (fuzzy sets) ([27]). Let *x* ∈ 𝒰 denote a measured value, where 𝒰 is the universe of discourse representing the full range of possible values for a given feature, such as expression levels or foldchanges. A feature *f* refers to a biological entity (e.g., gene, protein, or metabolite), and a sample corresponds to a single observation or experimental condition.

A fuzzy set *F* ⊆ 𝒰 is defined by a membership function (fuzzy value)

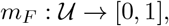

where *m*_*F*_ (*x*) quantifies the degree to which *x* belongs to the set *F* . Common membership functions include crisp (binary), triangular, trapezoidal, and Gaussian functions. These functions differ in how sharply they define category boundaries and how smoothly membership transitions occur.

A collection of fuzzy sets ℱ = {*F*_1_, …, *F*_*k*_} defined over the same universe forms a fuzzy concept. Fuzzification assigns each feature a membership vector, the fuzzy values

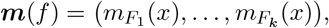

capturing partial membership across multiple linguistic labels such as ‘*low*’, ‘*medium*’, or ‘*high*’. This representation preserves uncertainty and allows fuzzy values to be propagated through downstream analyses.

### 2.2 Default Fuzzification

To define these fuzzy concepts automatically, one can make use of the law of large numbers. When enough measurements of the same kind are available, they typically follow a normal distribution.

For example, the foldchanges of all measured genes will be normally distributed around zero, as most genes are unchanged and will have a foldchange close to zero with deviation only because of noise, while the differential genes will be outliers in this normal distribution. This distribution can easily be fitted and used to define a Gaussian membership function for the ‘unchanged’ (‘◦’) fuzzy set. Trapezoidal memberships for up-regulated (’+’) and highly upregulated (’++’) are defined such that the intersect of the ‘unchanged’ and ‘+’ membership functions is at *σ* and of ‘+’ and ‘++’ at 3*σ* (see Figure 2 on the top left). The down-regulated fuzzy sets ‘−’ and ‘−−’ are defined similarly.

### 2.3 Fuzzifier^*^

In omics workflows, categorization is typically applied at the final stage of the analysis. Consequently, the analytical design choices that determine how and when categorization is introduced are often not examined explicitly, despite their substantial influence on the results.

Even in the simplest cases, there are two abstract analytical paths (see the black part of Figure 1 a). In the first path, analytical operations are performed in continuous space (A to B), and categorization is applied only to the final numerical result (B to E). In the second path, measurements are first mapped to fuzzy categories through fuzzification (A to D), and subsequent operations are carried out on these categorical representations (D to E).

**Figure 1:**
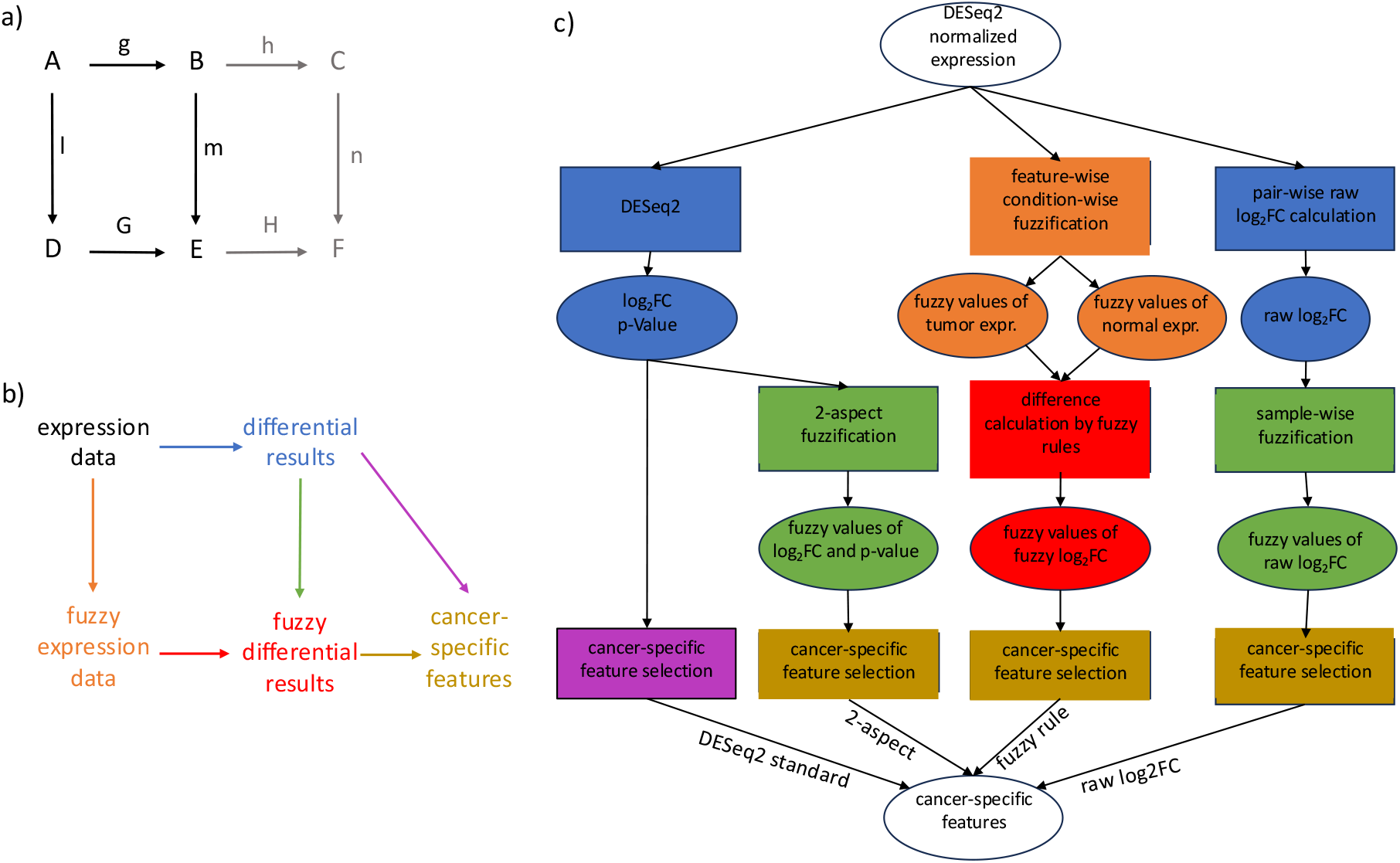
Overview of the analysis workflow. (a) shows the abstract commutative diagram, (b) the concrete instantiation for the identification of cancer-specific features, and (c) shows the Petri net workflow for different paths in the commutative diagram. Places and transitions in the Petri net are colored according to the arrows and data types in the commutative diagram.

Formally, these paths can be expressed as

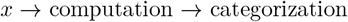

And

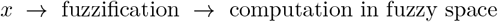

In both cases, the final outcome is a categorized representation defined over the same set of linguistic labels.

Because both paths terminate in the same categorical outputs, differences between the results can be attributed to the position of categorization within the path rather than to the choice of categories themselves. This abstraction makes it possible to evaluate fuzzification not as a replacement for existing analytical methods, but as an alternative placement of abstraction within the analysis pipeline.

As multiple computational steps are typically performed throughout a standard omics analysis pipeline, different combinations of calculations and categorization strategies can be applied (shown in the gray part of Figure 1, leading, for example, to F). This flexibility enables systematic evaluation of alternative analytical designs and allows the influence of both the choice and the position of categorization on the final results to be assessed.

### 2.4 Approaches for Fuzzy Differential Analysis

In general, defining cancer-specific miRNAs through differential analysis involves two steps: first, calculating differential expression results from the expression data, and second, determining which miRNAs are cancer-specific. Figure 1 b) shows the different ways to integrate fuzzification into the workflow. Either the expression data is fuzzified directly, and a fuzzy method is used to generate the fuzzy differential results, or the differential results are calculated from the normal expression values and then fuzzified. The approaches for these different paths in the commutative diagram are shown in more detail in Figure 1 c).

#### 2.4.1 DESeq2 standard

The standard DESeq2 approach does not use any fuzzification. The results of DESeq2 are simply filtered for miRNAs with an absolute log2 foldchange above 2 and corrected *p*-value below 10^−3^.

#### 2.4.2 DESeq2 2-aspect

A slight variant of the standard DESeq2 approach is to avoid hard thresholds by fuzzifying both log2 foldchanges and corrected p-values. The default fuzzification can be applied to the log2 foldchanges calculated by DESeq2 to fuzzify the log2 foldchanges. As the (corrected) *p*-values are not expected to be normally distributed, trapezoidal membership functions are defined over the −*log*_10_(*p* − *value*) to define different magnitudes of significance (from insignificant ‘◦’ to highly significant ‘^****^’). Intuitively, this corresponds to segmenting the volcano plot into different categories of effect size strength and significance (Figure 2).

**Figure 2:**
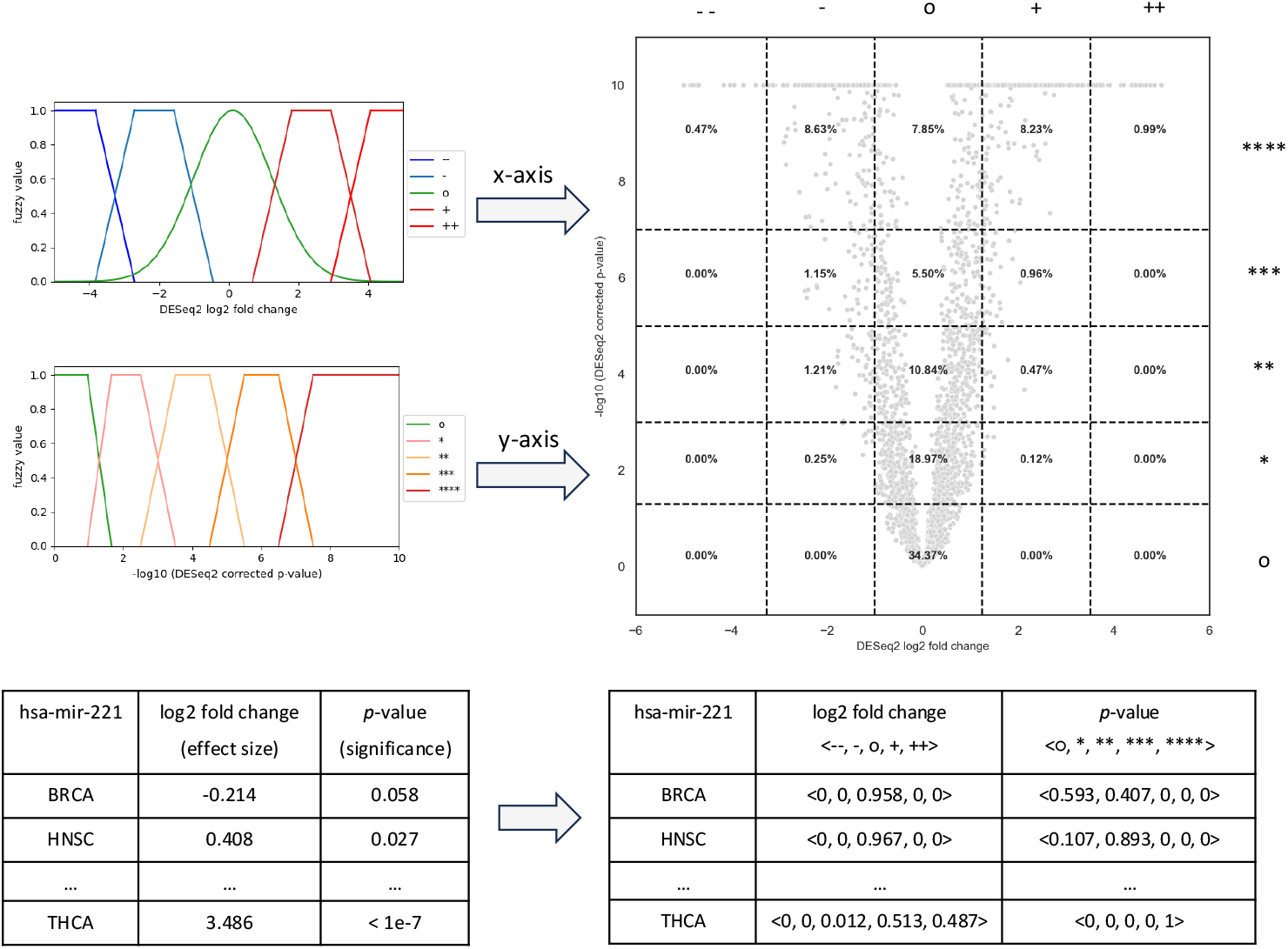
Fuzzy concept and volcano plot for log2 foldchange and negative log10-transformed corrected p-values from the DESeq2 statistical test. The horizontal and vertical dashed lines in the volcano plot represent the intersections between membership functions of neighboring fuzzy sets. The numbers in the subareas are the proportion of data points therein. For the fold-changes the default fuzzification is used, while trapezoidal membership functions are used for the ‘−*log*_10_(*corrected p* − *values*). Five fuzzy sets were defined both for DESeq2 log2 foldchange (‘−−’, ‘−’, ‘◦’, ‘+’, ‘++’) and the corrected *p*-values (’◦’, ‘^*^’, ‘^**^’, ‘^***^’, ‘^****^’). The raw and fuzzy values of one example miRNA hsa-mir-221 are shown in the tables under the figure. For each raw value, exactly one fuzzy value per fuzzy set is derived, resulting in a vector of fuzzy values with the same length as the number of fuzzy sets.

A miRNA is cancer-specific if it has either membership 1 in ‘−’/’+’ or non-zero membership in ‘−−’/’++’, respectively. Additionally, it has to have non-zero membership in ‘^****^’.

#### 2.4.3 Raw log2 foldchange (raw log2FC)

Another possibility to fuzzify the differential results is to first calculate for each pair of tumor and normal samples the log2 foldchange. As most miRNAs are expected to be unchanged, one can assume that this yields a normal distribution of log2 foldchanges that can be used for the default fuzzification. MiRNAs with more extreme log2 foldchanges will be in the ‘+’/’++’ or ‘−’/’−−’ categories.

The resulting fuzzy values are combined by averaging the fuzzy values over all tumor-normal sample pairs in each cancer type. A miRNA is cancer-specific in one tumor type if the sum of its average fuzzy value in ‘−∞’, ‘−−’ and ‘−’ (for down-regulated miRNAs) or ‘+∞’, ‘++’ and ‘+’ (for up-regulated miRNAs) exceeds a threshold of 0.5 and at least 60% of all tumor-normal sample pairs had their maximal membership in any of these three fuzzy sets.

In our analysis, the fuzzification was performed on the foldchanges between cancer and normal samples per patient to derive fuzzy membership values for each feature compared to the foldchange background of the other features. In the absence of explicit sample matching, samples can be paired combinatorially or according to experimental design, allowing the foldchange–based fuzzy interpretation to be extended to more general study setups.

#### 2.4.4 Fuzzy rule

Instead of fuzzifying the differential results, it is also possible to fuzzify the expression directly and then derive fuzzified differential results by a method that works on fuzzy values. To derive a suitable fuzzy concept, the default fuzzification is applied to the expression values of the normal samples (of a given cancer type). This is a typical measurement of replicates with a Gaussian error distribution, as they should only differ by noise. The derived fuzzy concept is then applied to both the tumor and normal samples.

To derive fuzzy differential results for one miRNA, Fuzzy Logic between the expression fuzzy values is used to derive differential classes. The tumor and normal fuzzy values of a sample pair are combined using vector multiplication, yielding fuzzy values for each combination of fuzzy sets in the tumor and normal sample. Table 1 shows the fuzzy rule matrix that maps each combination of tumor and normal fuzzy sets to a new differential fuzzy set. E.g., when both tumor and normal expression had membership 1 in ‘*NO*’, the differential fuzzy value will be membership 1 in ‘*NA*’. To reduce the complexity of the rules in this table, the memberships of ‘*low*’ and ‘*LOW*’ as well as ‘*high*’ and ‘*HIGH*’ are summed before the matrix multiplication. This operation is valid due to the additivity of fuzzification.

**Table 1:** Fuzzy rules for deriving the difference between fuzzy values of tumor (row) and normal (column) expression. Both fuzzy sets on either side of the fitted normal distribution are merged by adding the corresponding fuzzy values. Eight fuzzy sets, including three label fuzzy sets for ‘*NA*’ and ‘*±*∞’, are derived from the product matrix of vector multiplication.

|  | *NO* | *low+LOW* | *MEDIUM* | *high+HIGH* |
| --- | --- | --- | --- | --- |
| *NO* | NA | — | — — | — ∞ |
| *low+LOW* | + | o | — | — — |
| *MEDIUM* | ++ | + | o | — |
| *high+HIGH* | $+\infty$ | ++ | + | o |

And to derive the final differential fuzzy values, all entries in the rule matrix that lead to the same differential fuzzy set will be summed up. The final result of this combination is a membership vector of five levels of up-and down-regulation from ‘−−’ to ‘++’ and three extra fuzzy sets for labeling specific values ‘*NA*’ and ‘±∞’.

Cancer-specific miRNAs are identified in a similar way to the raw log2FC fuzzification. MiRNAs with a sum of average membership in fuzzy sets ‘−∞’, ‘−−’ and ‘−’ or ‘+∞’, ‘++’ and ‘+’ of at least 0.75 and that had their maximal membership in any of the three selected fuzzy sets in at least 75% of samples are defined to be cancer-specific.

### 2.5 Data acquisition and preprocessing

Raw counts of high-throughput miRNA-seq data from TCGA (The Cancer Genome Atlas Project, [2]) are downloaded for fuzzification. Since the final goal is to perform differential expression analysis using fuzzy logic, pairs of primary tumor and normal solid tissue samples from the same patient are built. Nine tumor types containing at least 30 such tumor-normal sample pairs are filtered to ensure sufficient number of replicates per tumor type for gaussian errors among replicates. The 1881 miRNAs are filtered according to their expression in primary tumor or solid normal tissue samples collected from the nine tumor types, namely 18 subconditions. They are expected either to be expressed in at least 50% of samples in all 18 subconditions or expressed in over 70% of samples in only one subcondition and less than 10% in all others. This results in 359 remaining miRNAs, where only hsa-mir-4686 is almost exclusively expressed in solid normal tissue samples from LIHC (liver hepatocellular carcinoma). The raw counts are normalized using DESeq2 [10], followed by log2-transform. Pair-wise log2 foldchanges are then calculated for each tumor-normal sample pair. Each comparison between primary tumor and solid normal tissue samples per tumor type is regarded here as one condition, which corresponds to the nine selected tumor types.

The cancer miRNA census (CMC, [22]) is used as a benchmark for the identified cancer-specific miRNAs. The study includes 634 miRNAs in different tumor types including TCGA, which are ranked by multiple criterion regarding databases, consistency, hallmarks, mutations and KEGG pathways. The miRNAs are assigned into two groups according to this ranking, namely CMC and non-CMC miRNAs, with a score threshold of 3.0.

## 3 Results

We applied the *Fuzzifier*^*\**^ framework to TCGA miRNA-seq data to systematically evaluate how different placements of categorization/fuzzification within the analysis pipeline affect differential feature detection. Across multiple analytical paths, *Fuzzifier*^*\**^ consistently recovered known cancer-associated miRNAs while also identifying novel, context-specific candidates that were missed by conventional threshold-based approaches.

There are 427 cancer-specific miRNAs *<miRNA, tumor type>* identified by either of the four methods, and 392 of them are validated by CMC with a score cutoff of 3.0. Figure 3 (a) shows the overlaps of these cancer-specific miRNAs between the different approaches. There are 49 cancer-specific miRNAs that were identified by all four methods, and quite a high proportion of them are also described by CMC. Around 37.5% of the cancer-specific miRNAs for both DESeq2-based methods are supported by CMC, while this percentage is higher for the fuzzy-based methods, raw log2FC and fuzzy rule, at over 50% of the identified cancer-specific miRNAs.

**Figure 3:**
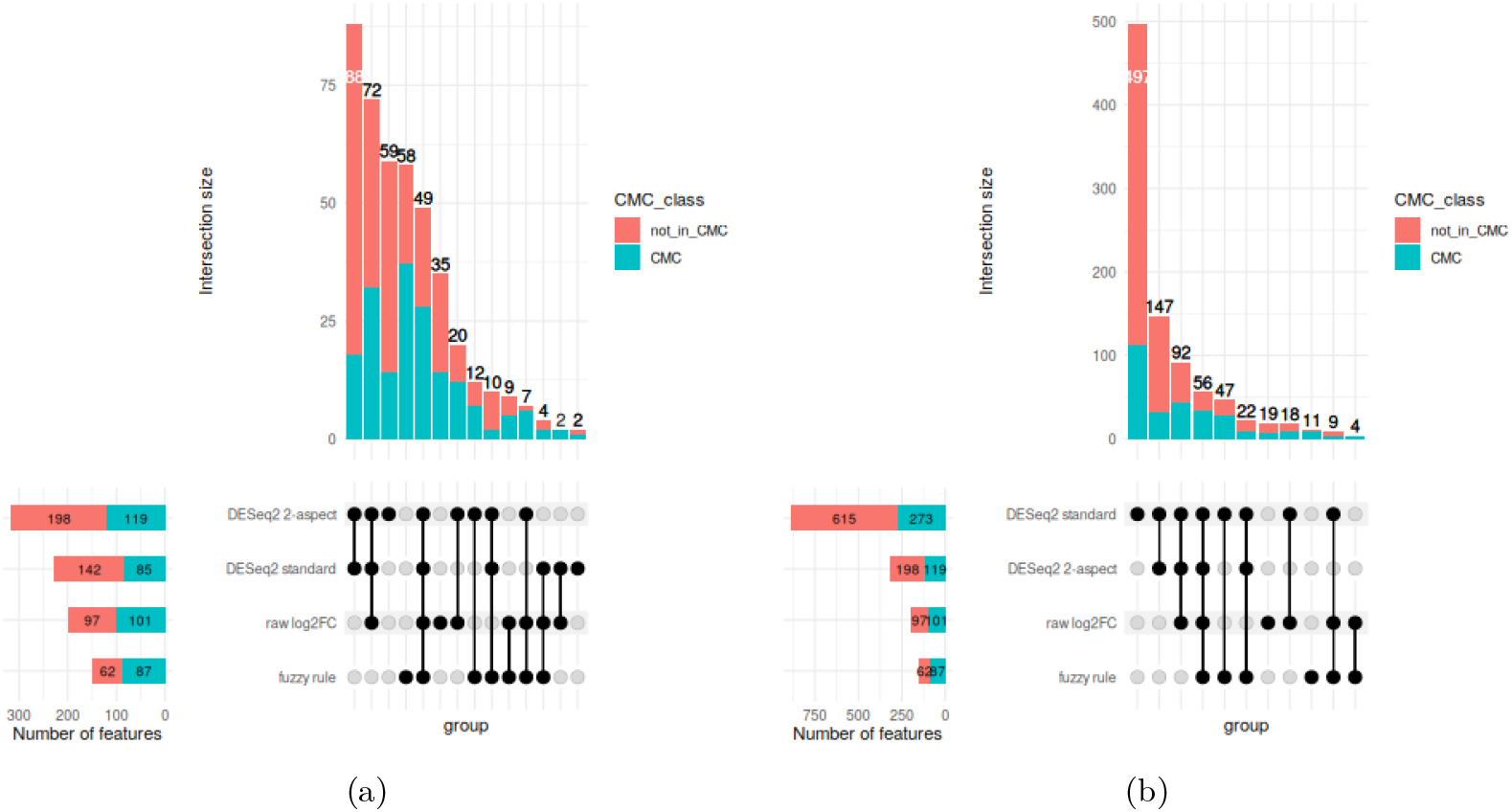
Upset plot for all identified cancer-specific miRNAs for DESeq standard with strict (a) and standard (b) thresholds. A miRNA with a CMC score of at least 3.0 is regarded as a CMC miRNA.

Fuzzy rule and raw log2FC, the two approaches that use fuzzification early in the pipeline, share only a small portion of identified cancer-specific miRNAs (69 of 198/148 for raw log2FC/fuzzy rule, respectively). This is expected as they require the miRNAs to show quite different behavior. While a miRNA has to have large enough changes of the pairwise log2 fold-changes in enough samples for raw log2FC, fuzzy rule requires a large enough difference in the distribution of the expression values.

Fuzzy rule and DESeq2 2-aspect have the highest number of cancer-specific miRNAs that are identified by no other method. The proportion of miRNAs in this subset that were also found by CMC is much higher for fuzzy rule than for DESeq2 2-aspect, which could indicate that the cancer-specific miRNAs identified by fuzzy rule might be more reliable.

DESeq2 2-aspect fuzzification finds the largest number among all four methods, followed by the DESeq2 standard method, with a total number of 227 identified cancer-specific miRNAs. The results of these two approaches overlap strongly, with an intersection of 219 cancer-specific miRNAs. This is expected as the approaches differ only in the way the cancer-specific miRNAs are selected. DESeq2 standard uses fixed cutoffs of 2 for foldchange and 10^−3^ for p-value, while DESeq2 2-aspect uses fuzzified versions of these values, which are then filtered to derive the binary cancer-specific or not categorization. This corresponds to a transformation of the cutoffs (approximately 1.6 for absolute log2 foldchange and 10^−6.5^ for corrected *p*-value), so that the main difference of these two approaches is that DESeq2 standard uses completely arbitrary thresholds, while the thresholds are at least partly derived from the underlying distribution (using the default fuzzification) in the DESeq2 2-aspect approach.

Here, we used quite strict thresholds for DESeq2 standard, as we mainly want to find the clear cancer-specific miRNAs. However, if traditional thresholds (1 for absolute log2 fold-change and 0.05 for corrected *p*-values) are applied (see Figure 3 (b)), 888 cancer-specific miRNAs are found, for which the DESeq2 2-aspect fuzzification results are a complete subset. Approximately 30.7% of these cancer-specific miRNAs were also described to be cancer-specific by CMC. Most miRNAs identified by only a single method were also detected by the DESeq2 standard approach using conventional thresholds, thereby increasing confidence that cancer-specific miRNAs uniquely identified by individual methods are nonetheless biologically relevant. The number of common cases identified by all four methods increases only slightly to 56.

We selected example miRNAs that were identified by only one of the approaches (DESeq2 standard and DESeq2 2-aspect, fuzzy rule or raw log2FC) or by all approaches and visualized the distribution of the expression values for the tumor and normal samples (see Figure 4). Hsamir-34a (a) and hsa-mir-221 (b) were found to be cancer-specific for thyroid carcinoma (THCA) by all four approaches, and both are also validated by CMC. The distribution of the normal and tumor expression values is both relatively compact and nicely separated from each other. In addition to the support by CMC, it has also been shown that hsa-mir-221 is overexpressed in thyroid papillary carcinoma and involved in cell cycle regulation by inhibiting CDKN1B expression [23]. Similarly, [11] showed that hsamir-34a inhibits GAS1 expression as its direct target, leading to altered functionality of the PI3K/Akt/Bad pathway and promoting tumorigenesis.

**Figure 4:**
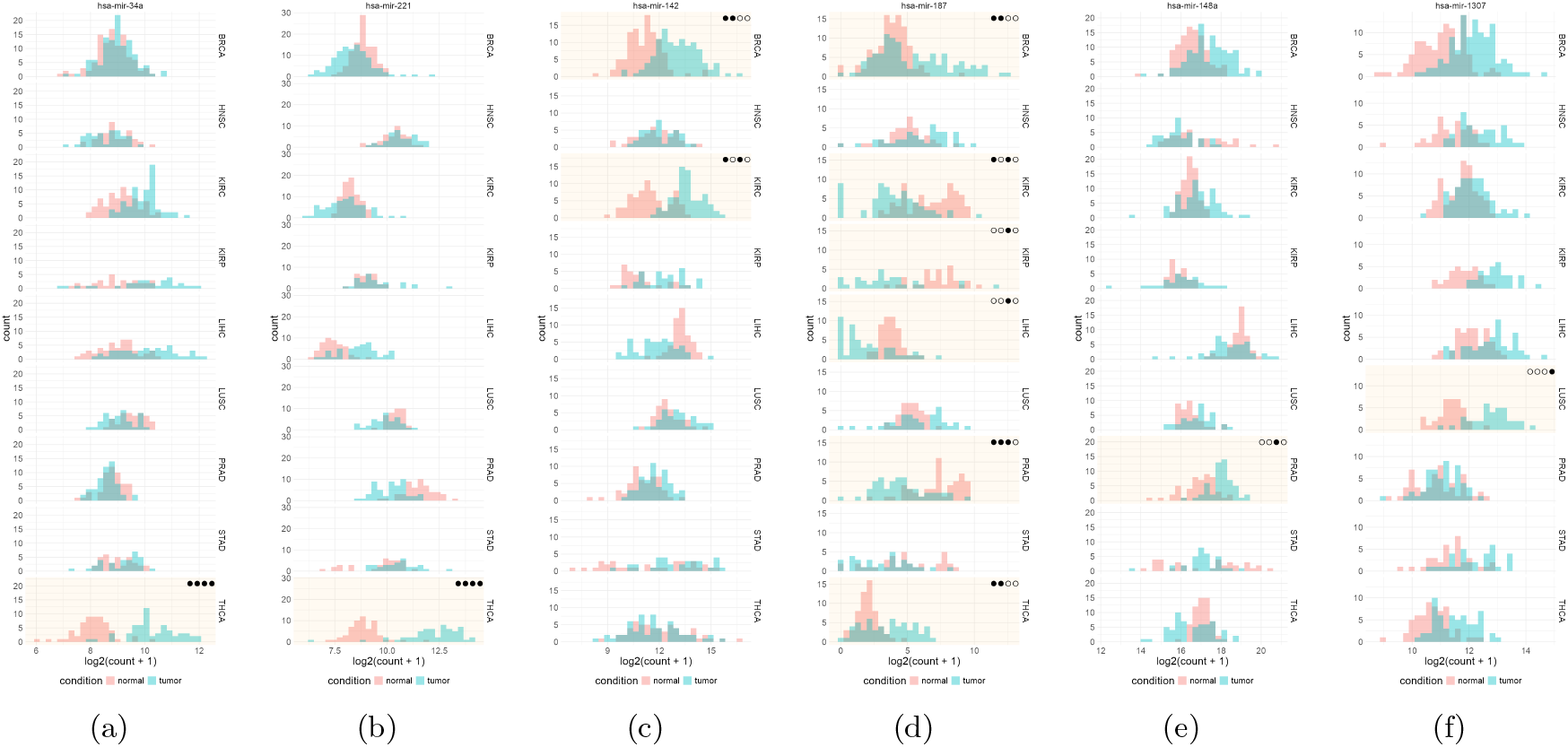
Histograms of tumor (blue) and normal (pink) expression distribution in the nine selected tumor types, for example, cancer-specific miRNAs. The miRNAs were selected to be identified by all four methods (a, b), only by DESeq2 2-aspect fuzzification or DESeq2 standard method (c, d), only by fuzzy rule (e), and only by raw log2FC (f). The four dots on the upper right corner of each shaded histogram represent the method(s) by which the cancer-specific miRNA is identified (order: DESeq2 2-aspect, DESeq2 standard method, raw log2FC and fuzzy rule).

The distributions of the two examples that were found only by the two DESeq2-based methods (hsa-mir-142 (c) and hsa-mir-187 (d), both in BRCA) overlap nearly completely. While there is a shift towards upregulation in the tumor samples, it is not pronounced enough to be found by the other approaches. For hsa-mir-187, many of the tumor samples are even at the lower end of the normal sample distribution, while other tumor samples are clearly upregulated. This could indicate that this miRNA is only upregulated in a subset of patients, and could even be a subtype marker. Indeed, several studies have reported this miRNA as a biomarker associated with poor prognosis in breast cancer [13, 1]. While this is of course interesting, it does not correspond to our goal of finding miRNAs that are cancer-specific in most patients.

Hsa-mir-148a is an example of a miRNA that was only found by the raw log2FC approach to be specific for prostate cancer (PRAD). The expression distributions are shifted but overlap substantially. However, it is not clear which values are paired in the expression distributions, and as raw log2FC analyses the paired foldchange distribution and finds this miRNA, it is likely that most paired foldchanges are upregulated. [3] and described that hsa-mir-148a is a biomarker to differentiate malignant prostate cancer from benign lesions.

Fuzzy rule was the only approach that identified hsa-mir-1307 as specific for lung squamous cancer (LUSC). The expression distributions are separated, but maybe not as clearly as for the examples that were identified by all methods. [4] found that it upregulates the MAPK pathway, leading to proliferation in lung cancer.

Figure 5 shows examples for the analysis using conventional thresholds for DESeq2 standard. The miRNAs that were found only by DESeq2 standard with conventional thresholds do not show a clear separation of the normal and tumor expression distributions, but look more like the example in Figure 5(a) where both distributions overlap nearly completely with only a slight shift and some outliers. Here, 2-aspect would require somewhat stronger shifts even for the strong DEseq2 p-value, and thus does not find this miRNA as cancer-specific. While most of the miRNAs that were found uniquely by fuzzy rule or raw log2FC are also found by DESeq2 with conventional thresholds, there are still examples only found via other paths: Figure 5(b) shows an example of a miRNA that was only found by fuzzy rule, but no other method. The two distributions are clearly separated, so that one would accept this miRNA as being differential. For the examples that were only found by raw log2FC the separation of the distributions is not that clear. Hsa-mir-148a, that was already our example miRNA only found by raw log2FC in Figure 4(e) is also not identified as differential by DESeq2 with conventional thresholds, even though it is a known prostate cancer-specific gene. There is also a shift between the distributions in PRAD comparable to the shift observed for BRCA and HNSC which are both differential according to DESeq2.

**Figure 5:**
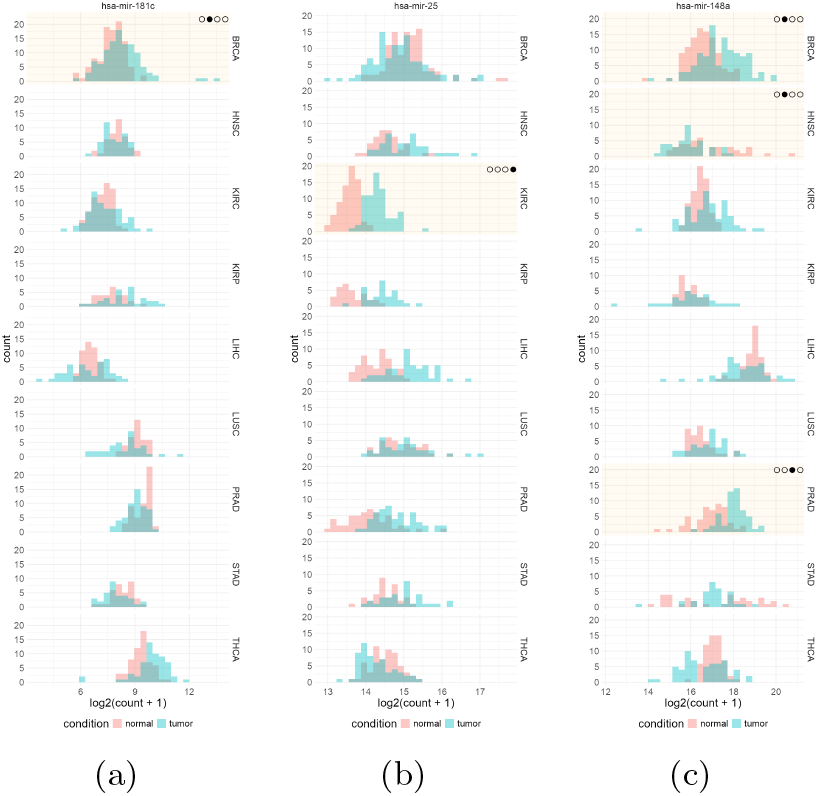
Histograms of tumor (blue) and normal (pink) expression distribution in the nine selected tumor types for the example cancer-specific miRNAs. The miRNAs were selected to be identified only by DESeq2 standard method with conventional thresholds (a), only by fuzzy rule fuzzification (b) and only by raw log2FC fuzzification (c) (order for the dots: DESeq2 2-aspect, DESeq2 standard method, raw log2FC and fuzzy rule).

## 4 Discussion

Decision-making in data analysis remains inherently challenging, particularly when complex statistical transformations and modeling techniques are involved. While existing tools provide powerful analytical capabilities, the responsibility for interpreting results and selecting meaningful outcomes ultimately lies with the user. In this context, our approach aims to reduce subjective uncertainty by actively informing and guiding users toward more trustworthy and interpretable results. Rather than enforcing a single analytical path, it acknowledges that multiple valid interpretations of the same data may coexist and can be meaningfully integrated into a holistic outcome. At the same time, different criteria may be more relevant for specific analyses and can be explicitly assessed by introducing additional paths within the framework.

The use of fuzzification represents a key contribution of this work, as it enables value categorization based on the underlying data distribution in a manner that more closely resembles human perception. With sufficiently large sample sizes (such as those available from high-throughput sequencing data generated by TCGA), the law of large numbers permits reliable fitting of normal distributions to real-valued expression data. This distribution-aware representation provides a foundation for subsequent fuzzy-rule-based processing, enabling analyses that go beyond conventional threshold-based decision-making.

We show that fuzzification can be integrated at multiple steps into the analysis pipeline. As the different approaches use different properties of differential features, such as how well the expression distributions were separated (fuzzy rule) or whether most samples show a large enough foldchange in the paired setup (raw log2FC), it is not surprising that their results differ. Fuzzifer^*^ allows for exploring these different paths in the commutative diagram. How the results of these paths are integrated depends on the use case: when possible, drug targets should be identified; one is interested only in the clearest cases, which are found by all paths. But if the resulting differential features should be used for further analyses, like e.g. understanding pathways relevant in cancer, one would want to be more sensitive and would maybe even include all features that were identified by any path.

In our use case, we observed that standard approaches yield very long result lists with conventional thresholds, and that even for quite strict thresholds, there are examples that are not that clearly differential. For advanced applications, statistical significance alone is frequently insufficient; for example, a clear separation of samples in raw expression space or foldchange space can be critical. It is therefore important to incorporate additional criteria to support more informed and context-dependent decision-making. Consequently, multiple analytical approaches should often be applied, extending beyond traditional conventions and the strict enforcement of underlying assumptions. Assumptions, such as specific distributions, are frequently required on the data but rarely tested in practice. As a result, both the data and the outcomes can be affected, leading to significant results that differ substantially from what would be expected.

We have shown that *Fuzzifier*^*\**^ is a novel, generally applicable categorization tool, which allows us to define additional analysis paths for given analysis pipelines via systematically adding explicit categorizations for intermediate results. The respective categorizations rely on user-defined data-driven data modeling (fuzzification, e.g., with the *Fuzzifier* tool), enabling the generation of additional ‘views’ on the data.

The quite different results of the investigated paths can be interpreted in more conservative (intersections) or more sensitive (unions) ways to increase the ‘degree of belief’ in the accepted predictions. The views and *Fuzzifier* reports provide additional (semi-)quantitative measures and visualizations, which increase or decrease the evidence derived from the data at hand for the claimed hypotheses and findings.

Thus, *Fuzzifier*^*\**^ is a general method to obtain reproducible, robust, transparent and trusted findings. It is often not really possible to derive causal conclusions such as ‘miRNA x is differentially regulated in cancer c’ or ‘miRNA y is a diagnostic/prognostic marker for cancer (sub-)type z’ from any (likely not large enough and not targeted enough) study data. In order to conclude about causal relations with beyond mere correlations derived from the available study data, more and custom-designed perturbation experiments may be still be required, but *Fuzzifier*^*\**^ should be of help to best use the available data and prior knowledge and to reduce superfluous experiments by concentrating on the best hypotheses implied by the available data.

## 6 Author contributions statement

FO, CP: Conceptualization, Methodology, Data Curation, Software, Visualization, Investigation, Writing -Original Draft; ES: Visualization, Investigation, Supervision, Writing; RZ: Conceptualization, Writing, Supervision, Funding

## 7 Acknowledgments

Part of this work has been funded by DFG CRC1123 Atherosclerosis and the PhD program of the LSB.

